# Costing ‘the’ MTD … in 2-D

**DOI:** 10.1101/370817

**Authors:** David C. Norris

**Author notes:** Corresponding author: David C. Norris.

## Abstract

**Background:** I have previously evaluated the efficiency of one-size-fits-all dosing for single agents in oncology (Norris 2017b). By means of a generic argument based on an E_max_-type dose-response model, I showed that one-size-fits-all dosing may roughly halve a drug’s value to society. Since much of the past decade’s ‘innovation’ in oncology dose-finding methodology has involved the development of special methods for *combination therapies*, a generalization of my earlier investigations to *combination dosing* seems called-for.

**Methods:** Fundamental to my earlier work was the premise that *optimal dose* is a characteristic of each individual patient, distributed across the population like any other physiologic characteristic such as height. I generalize that principle here to the 2-dimensional setting of combination dosing with drugs A and B, using a copula to build a bivariate joint distribution of (MTD_*i,A*_, MTD_*i,B*_) from single-agent marginal densities of MTD_*i,A*_ and MTD_*i,B*_, and interpolating ‘toxicity isocontours’ in the (*a, b*)-plane between the respective monotherapy intercepts. Within this framework, three distinct notional toxicities are elaborated: one specific to drug A, a second specific to drug B, and a third ‘nonspecific’ toxicity clinically attributable to either drug. The dose-response model of (Norris 2017b) is also generalized to this 2-D scenario, with the addition of an interaction term to provide for a complementary effect from combination dosing. A population of 1,000 patients is simulated, and used as a basis to evaluate population-level efficacy of two pragmatic dose-finding designs: a dose-titration method that maximizes dose-intensity subject to tolerability, and the well-known POCRM method for 1-size-fits-all combination-dose finding. Hypothetical ‘oracular’ methods are also evaluated, to define theoretical upper limits of performance for individualized and 1-size-fits-all dosing respectively.

**Results:** In our simulation, pragmatic titration attains 89% efficiency relative to theoretically optimal individualized dosing, whereas POCRM attains only 55% efficiency. The passage from oracular individualized dosing to oracular 1-size-fits-all dosing incurs an efficiency loss of 33%, while the parallel passage (within the ‘pragmatic’ realm) from titration to POCRM incurs a loss of 38%.

**Conclusions:** In light of the 33% figure above, the greater part of POCRM’s 38% efficiency loss relative to titration appears attributable to POCRM’s 1-size-fits-all nature, rather than to any pragmatic difficulties it confronts. Thus, appeals to pragmatic considerations would seem neither to justify the decision to use 1-size-fits-all dose-finding designs, nor to excuse their inefficiencies

## INTRODUCTION

Until now, I have advanced my criticism of 1-size-fits-all dose-finding methods strictly in the realm of monotherapy. (Norris 2017a) advocated a philosophy of *adaptive dose individualization*, under the heading ‘dose titration algorithm tuning’ (DTAT), and illustrated this concretely through simulated neutrophil-nadir-targeted titration of a chemotherapy agent to an individualized ‘MTD_*i*_’ that varies from one patient to another. (Norris 2017b) examined ‘1-size-fits-all’ specifically as a dose-finding *constraint*—and estimated the social costs of this constraint as a function of CV(MTD_*i*_), the coefficient of variation of MTD_*i*_ in the population. (Norris 2017c) elaborated a new coherence principle in dose finding, *precautionary coherence* (PC), and employed it to motivate a pragmatic ‘3+3/PC’ dose-titration design that beats any conceivable 1-size-fits-all design on both ethical and utilitarian grounds. The present paper begins a process of extending these earlier arguments into the realm of *combination therapy*.

## TOXICITY ISOCONTOURS FOR 2-AGENT THERAPY

It has been recognized for at least a quarter-century (Korn and Simon 1993) that the combination-therapy setting spoils the determinacy of dosing identified on a maximum-tolerated-dose (MTD) heuristic. Formally speaking, in combination dosing with *n* agents, the set of tolerable doses will in general be bounded by an (*n −* 1)-dimensional manifold of just-barely-tolerable dose combinations. Specifically in the case of 2 agents A and B, this is a curve in the (*a × b*)-plane.

Like so many concepts in dose finding, the problem of *combination dose finding* yields up fresh conceptual clarity when we consider it from an individual-patient perspective. Considered as belonging to individual patients, the abovementioned curves arise naturally as isocontours of the individual’s disutility *U_i_*(*a, b*) for the joint toxicities of a dose. Operating on the MTD heuristic amounts to identifying a disutility 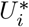 that defines a just-barely-tolerable dose for individual *i*, and seeking a combination dose somewhere on the *toxicity isocontour* 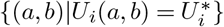.

It would introduce a distracting level of detail into the present discussion, however, to model such utilities explicitly here. Instead, I obtain individual-level toxicity isocontours in a manner that more directly generalizes my earlier single-agent methodological investigations (Norris 2017b). Specifically, I suppose that for each specific toxicity *tox*, the associated single-agent MTD’s 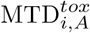 and 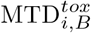 are marginally Gamma-distributed:

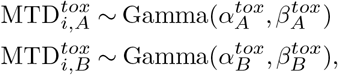

with their joint distribution constructable using the Clayton copula:

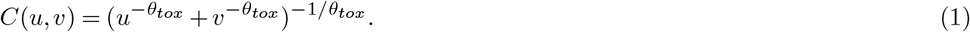

From this joint distribution, we may draw pairs 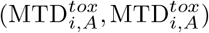 as in Figure 1. These yield the endpoints 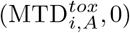 and 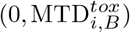 of the toxicity isocontours, which we interpolate in the (*a, b*)-plane using solutions of

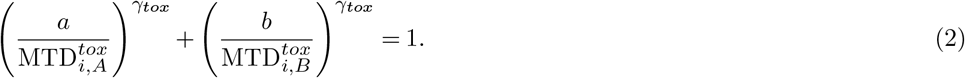

**Figure 1.**
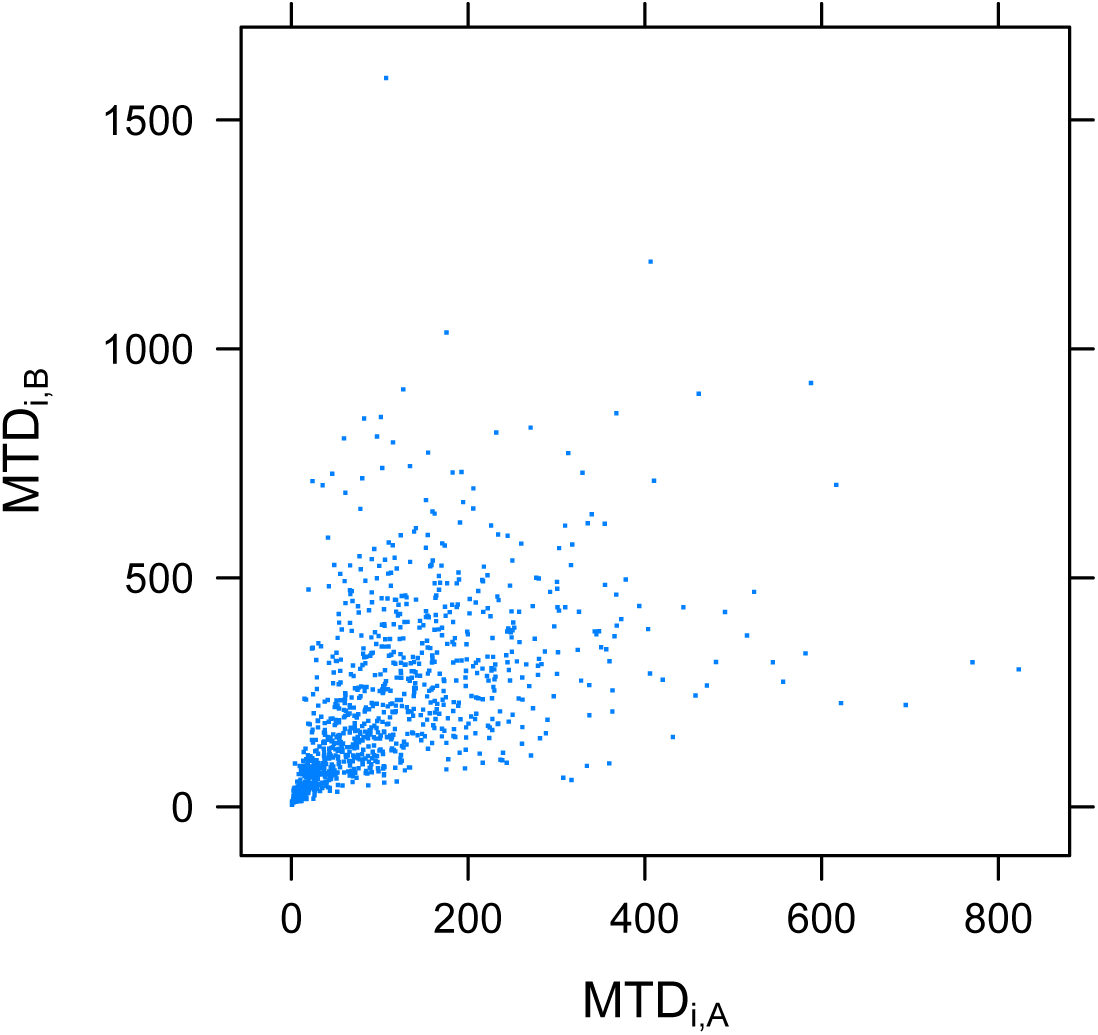
Joint density of *monotherapy* MDT_*i*_’s for 2 drugs, relative to a given toxicity, *tox*. The marginal distributions are of the Gamma type, with scale parameters 1*/β_A_* = 100 and 1*/β_B_* = 150, and coefficients of variation 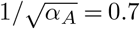 and 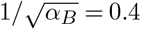.The joint density is constructed using the Clayton copula (Equation 1) with parameter *θ_tox_* = 2.

With *γ_tox_* ∈ (0, ∞), Equation 2 can model a range of toxicity interactions, as shown in Figure 2. At *γ_tox_* = 1, we obtain a straight line consistent with a purely additive toxicity, such as would occur when A and B produce the toxicity by a shared mechanism.^1^ When *γ_tox_ <* 1 we have by comparison a relatively synergistic (i.e., *more-than-additive*) toxicity, while for *γ_tox_ >* 1 drugs A and B produce the toxicity relatively independently. As *γ_tox_* → *∞*, the toxicities of A and B tend toward a complete independence.

**Figure 2.**
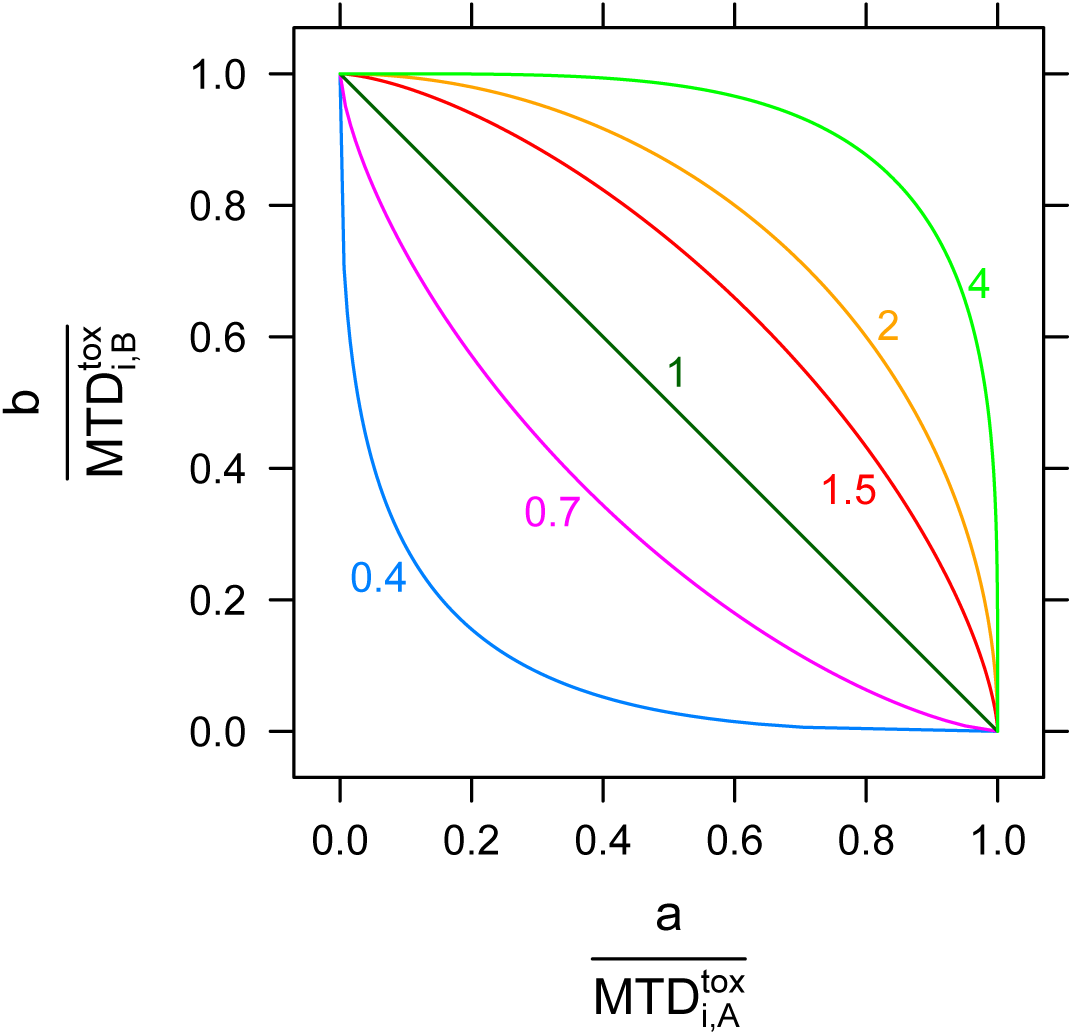
Toxicity isocontours generated by Equation 2 with *γ_tox_* ranging from 0.4 to 4.

## THREE TOXICITIES, WITH INTER-INDIVIDUAL HETEROGENEITY

A perennial failing in the biostatistical dose-finding literature is its abnegation of clinical knowledge and biological understanding of toxicity. With respect to *combination* dose finding, in particular, this manifests most conspicuously as the presumption that specific toxicities cannot be clinically attributed to one of the combination agents. (This literature does contain isolated forays into toxicity attribution; see e.g. (Jimenez, Tighiouart, and Gasparini 2018) and the sparse prior efforts surveyed therein.) To provide a concrete and clinically reasonable basis for considering toxicity attribution, I posit 3 notionally distinct toxicities: one specific to drug A, another to drug B, and a third ‘nonspecific’ toxicity clinically attributable to either A or B. See Table 1.

**Table 1.** Three notional toxicities differing in their clinical attributability to drugs A and B of our hypothetical combination therapy.

| Toxicity | Implicated Drug | Notional example drug |
| --- | --- | --- |
| Neutropenia | A | Cytotoxic chemotherapy |
| Diarrhea | B | Checkpoint inhibitor |
| Fatigue | A or B | Many |

To implement this concept within the toxicity-isocontour framework set forth above, we treat each toxicity as defining its own 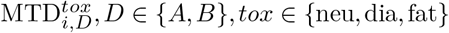, where (e.g.) the attributability of *neutropenia* and *diarrhea* to drugs A and B respectively emerges because

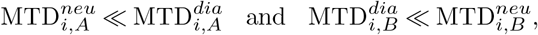

while the nonspecific nature of *fatigue* reflects that

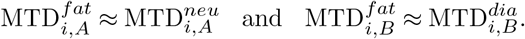

Indeed, if we render the doses of drugs A and B *commensurate* by rescaling them to unity in the ranges where their characteristic toxicities occur:

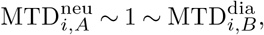

then the clinical logic of Table 1 translates roughly to

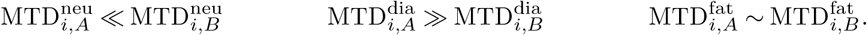

Assuming that the disutility of multiple distinct toxicities is determined by the disutility of the *worst* toxicity^2^, then these ideas lend themselves to straightforward treatment in a simple planar diagram. Each toxicity divides the (*A, B*) dose plane into two regions: tolerable and intolerable. The intersection of the tolerable regions with respect to each separate toxicity yields the set of dose combinations tolerable with respect to *all* toxicities. These tolerable regions are the white areas close to the origin in Figure 3.

**Figure 3.**
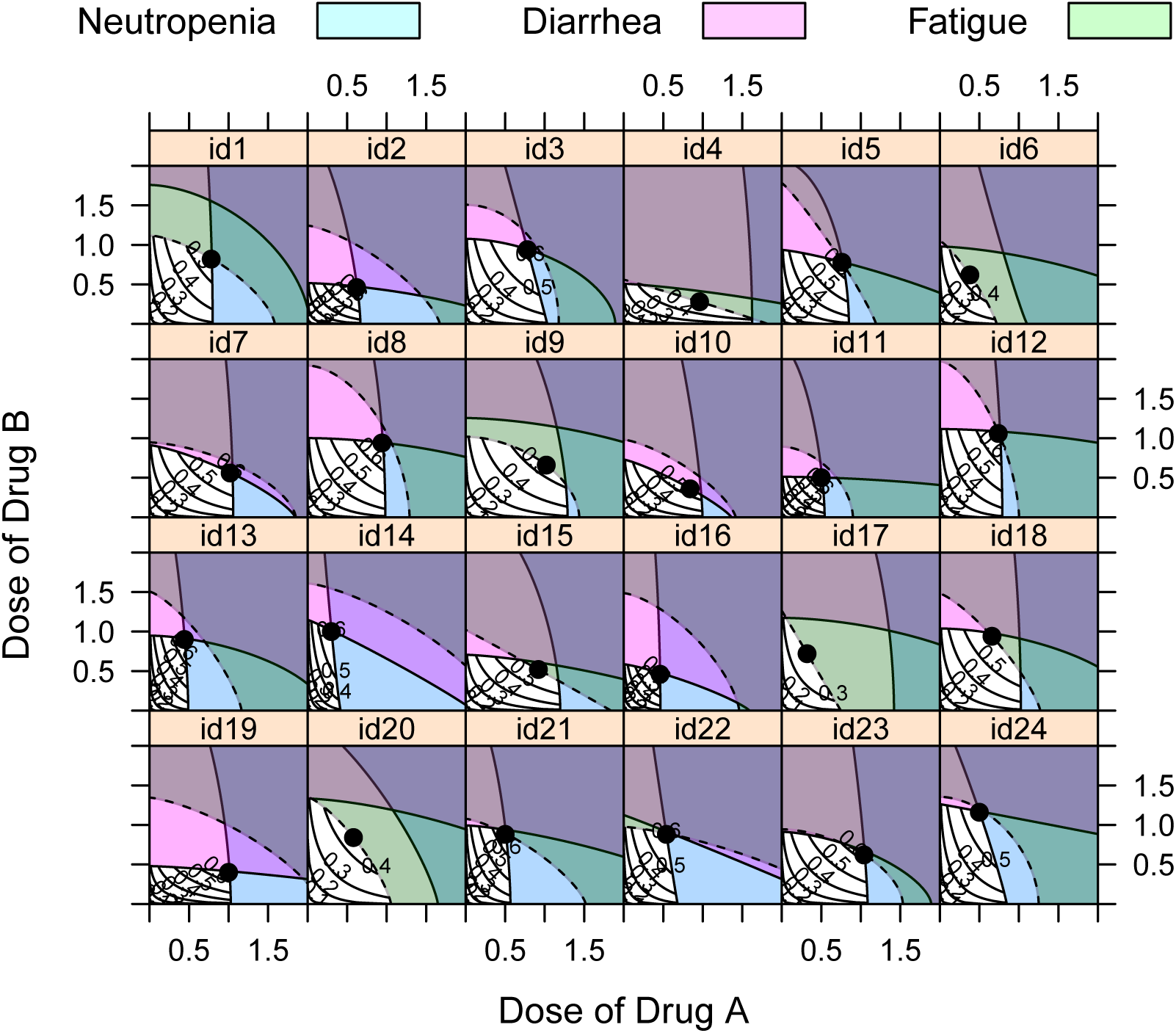
Heterogeneous combination-dosing constraints imposed by the 3 notional toxicities of Table 1, for 24 different patients. The regions of (*a, b*)-dose space rendered intolerable by the 3 toxicities are filled with distinct colors. The attributability of *neutropenia* and *diarrhea* to drugs A and B corresponds to the verticality and horizontality of the respective toxicity isocontours. Likewise, the inattributability of *fatigue* reflects that its isocontours (dashed) typically have intercepts on both A and B axes. Within the (white) *tolerable regions* are plotted *level curves* of the probability of remission *P_r_*(*a, b*) from Equation 3. The black dot in each panel shows the tolerable combination dose that yields optimal efficacy for each individual. For half of the individuals shown, the optimal dose occurs at the intersection of the neutropenia and diarrhea contours (look for the half-magenta, half-blue ‘bowties’), meaning that each of these toxicities is barely-tolerable at the optimal dose. For id1, it is neutropenia and fatigue which jointly constrain the optimal dose. For id18, the optimal dose produces dose-limiting diarrhea and fatigue. For id6, id9, id17 and id20, fatigue alone constrains the optimal dose, while for id10 only diarrhea constrains the optimal dose.

## DOSE-RESPONSE MODEL

The remission probability level curves in the tolerable regions of Figure 3 are derived from

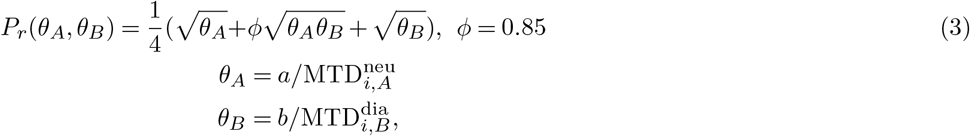

which generalizes the dose-response model of (Norris 2017b, Equation 1) to the case of 2-drug combination therapy.^3^ The choice of *φ* = 0.85 *>* 0 models a complementary (more-than-additive) effect such as we would typically seek from combination therapy.

## RELATIONS WITH KORN & SIMON’S ‘TOLERABLE-DOSE DIAGRAM’

Figure 3 bears a strong resemblance to the ‘tolerable-dose diagram’ which (Korn and Simon 1993) develop as graphical guidance to the design and conduct of combination dose-finding studies. The main difference conceptually between my Figure 3 and Korn & Simon’s Figures 2–6 is that the latter define ‘tolerability’ according to the population-level DLT *rates* that are typically targeted in 1-size-fits-all dose finding. Most of the guidance Korn & Simon articulate translates readily into a dose individualization setting, and indeed all of the core intuitions on which their paper depends seem to derive more directly and naturally from individual-level considerations. Thus, I regard the ‘toxicity isocontours’ of Figure 3 as achieving a *more faithful* representation of Korn & Simon’s original intuitions than could their own tolerable-dose diagram, burdened as it was with the DLT-rate conceptual baggage of 1-size-fits-all dose finding.

## THE COST OF THEORETICALLY OPTIMAL 1-SIZE-FITS-ALL DOSING

The scenario of Figure 3 provides a setting within which the performance of combination dose-finding methods can be examined and compared. Before undertaking (in the next section) to examine the performance of an *actual* 1-size-fits-all dose-finding method, I wish first to characterize the losses attributable to *1-size-fits-all dosing itself*, apart from the vagaries of any given method. In this section, therefore, we examine the performance of an ‘oracular’ 1-size-fits-all dosing regime that maximizes the population-rate of remission. As in (Norris 2017b), we define this as the single dose which, when given to all patients who can tolerate it, achieves the highest population rate of remission.

By this definition, the optimal 1-size-fits-all dose for 1000 simulated patients (of whom Figure 3 shows just the first 24) turns out to be (*a* = 0.44*, b* = 0.42). At this dose, 9.5% of patients have a DLT, and the population rate of remission is 0.396; see Figure 4. Relative to the population rate of 0.586 achieved under ‘oracular’ individualized dosing, this represents a 32.5% loss of efficacy.

**Figure 4.**
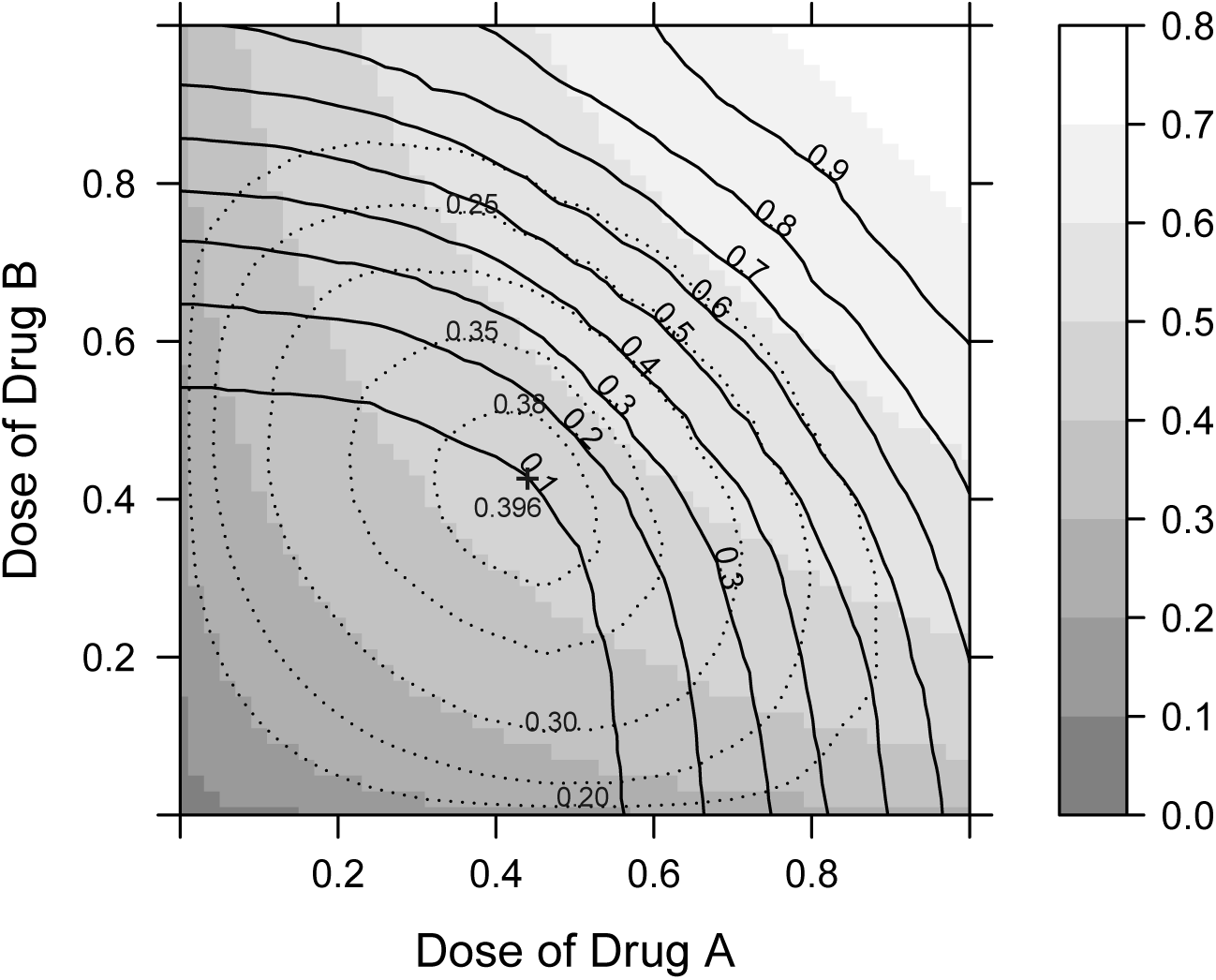
Contours of the DLT rate in the (*a, b*)-plane, superimposed on population rates of remission. The grayscale shows remission probabilities as driven by the dose-response model in Equation 3, irrespective of dose tolerability. The dotted contours show remission rates achieved under 1-size-fits-all dosing, where patients who cannot tolerate ‘the’ dose receive no treatment. The peak remission rate of 0.396 attained at the optimal 1-size-fits-all dose is marked with a ‘+’.

The shared toxicity introduced here complicates a direct comparison with the monotherapy setting of (Norris 2017b). Nevertheless, with all 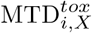 in our simulation sharing the same shape parameter *α* = 8, I find it hard to resist substituting this into Equation 3 of that previous paper. This yields a value of *P̂r* (*α*) = 0.346, corresponding to an efficacy loss of 30.9% relative to the population rate of 0.5 which (Norris 2017b) posited for optimal, individualized dosing.

The close proximity of these efficacy losses need not surprise us too much, especially in light of the generic set-up of (Norris 2017b) (requiring no special appeal to a single toxicity), and the robustness of its findings to the several sensitivity analyses performed therein. As the next section will demonstrate, the transition to ‘higher dimensions’ in dose finding most profoundly affects the very issue that in this section we have assumed away by taking our optima for granted.

## PERFORMANCE OF POCRM

To implement the practical search for a 1-size-fits-all dose, we select the well-known POCRM approach (Wages, Conaway, and O’Quigley 2011). This method builds on the CRM of (O’Quigley, Pepe, and Fisher 1990), far and away the most popular model-based method for 1-size-fits-all dose finding in monotherapy (Iasonos and O’Quigley 2014). The novel aspect of POCRM is its employment of partial ordering (PO) notions to address the *loss of monotonicity* which characterizes dose-response in the combination-therapy setting.

I implement POCRM on the 4× 4 grid of doses {*d*_1_, … *d*_16_} shown in Figure 5, following the lead of (Wages, Conaway, and O’Quigley 2011) as to the grid dimensions. In order to reflect the intention to seek truly *combination* dosing, I however drop doses *d*_4_ and *d*_13_ from further consideration.^4^ Also following (Wages, Conaway, and O’Quigley 2011), I posit 3 possible toxicity orderings (models) over these dose levels: {*m*_1_*, m*_2_*, m*_3_}. The first, *m*_1_, is the true ordering, derived from the toxicity rates observed in our 1000 simulated patients. The latter two, *m*_2_ and *m*_3_, are obtained by systematically distorting these probabilities. See Figure 6 and Table 2.

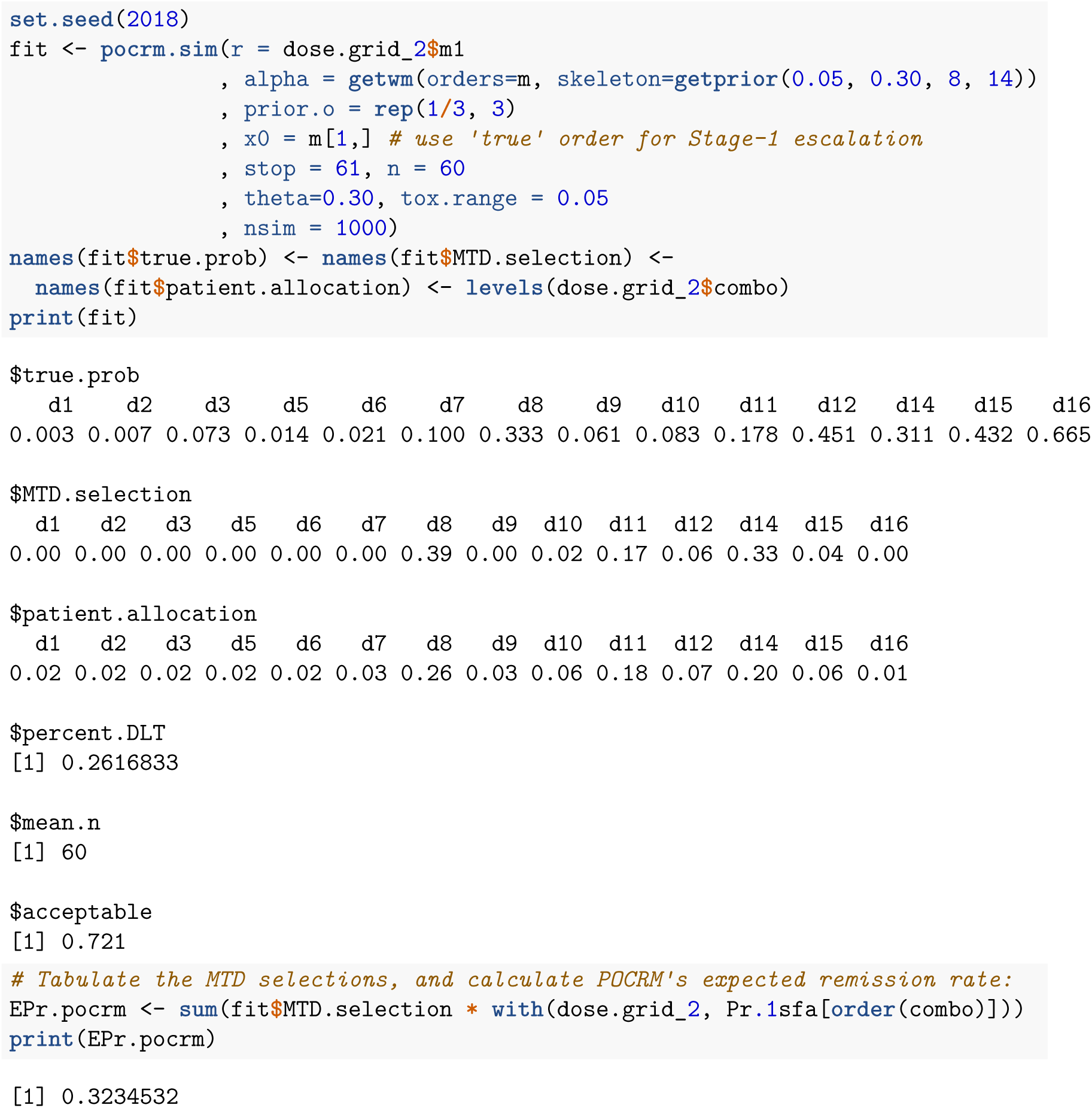

**Figure 5.**
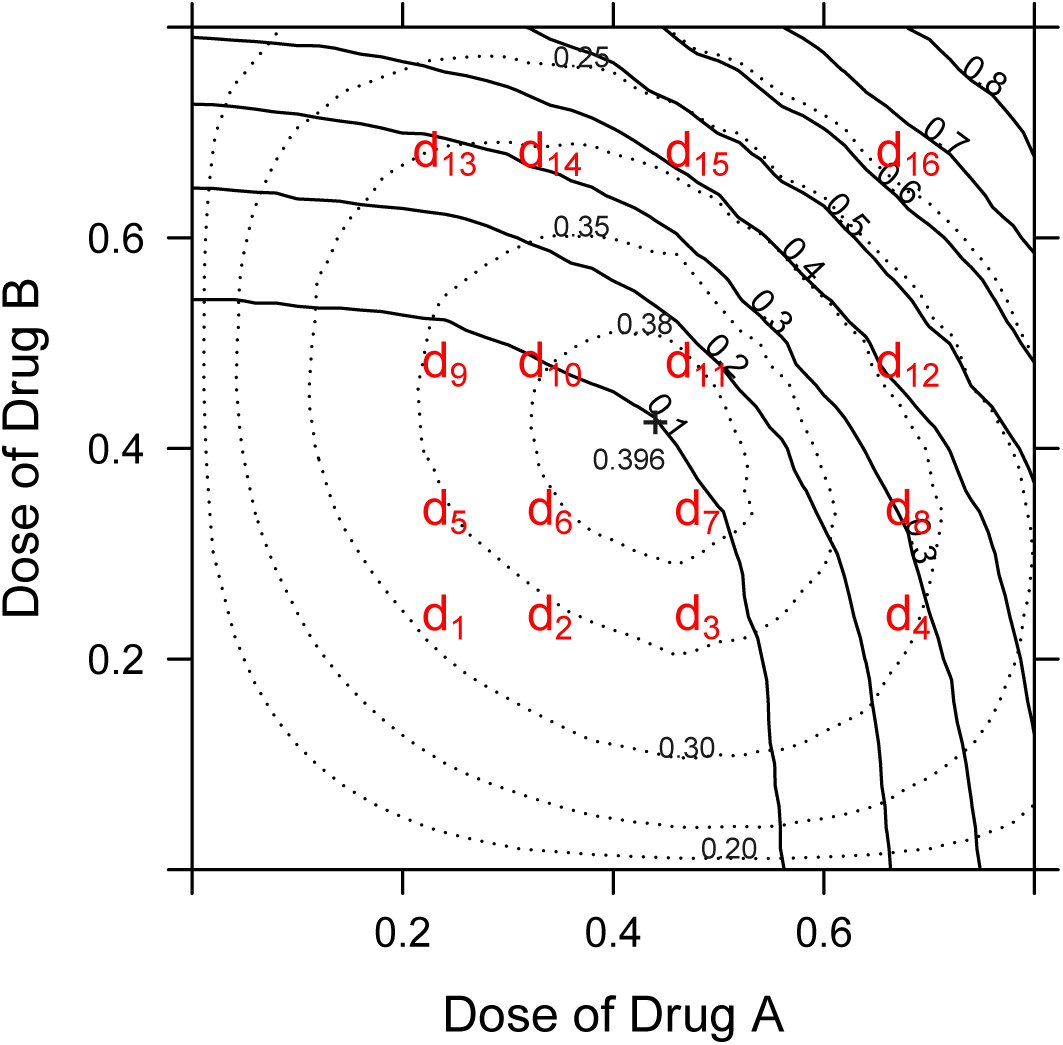
The geometrically-spaced grid of combination doses {*d*_1_, …*, d*_16_} over which practical search is conducted for optimal dose. The grid is shown superposed on a portion of Figure 4.

**Figure 6.**
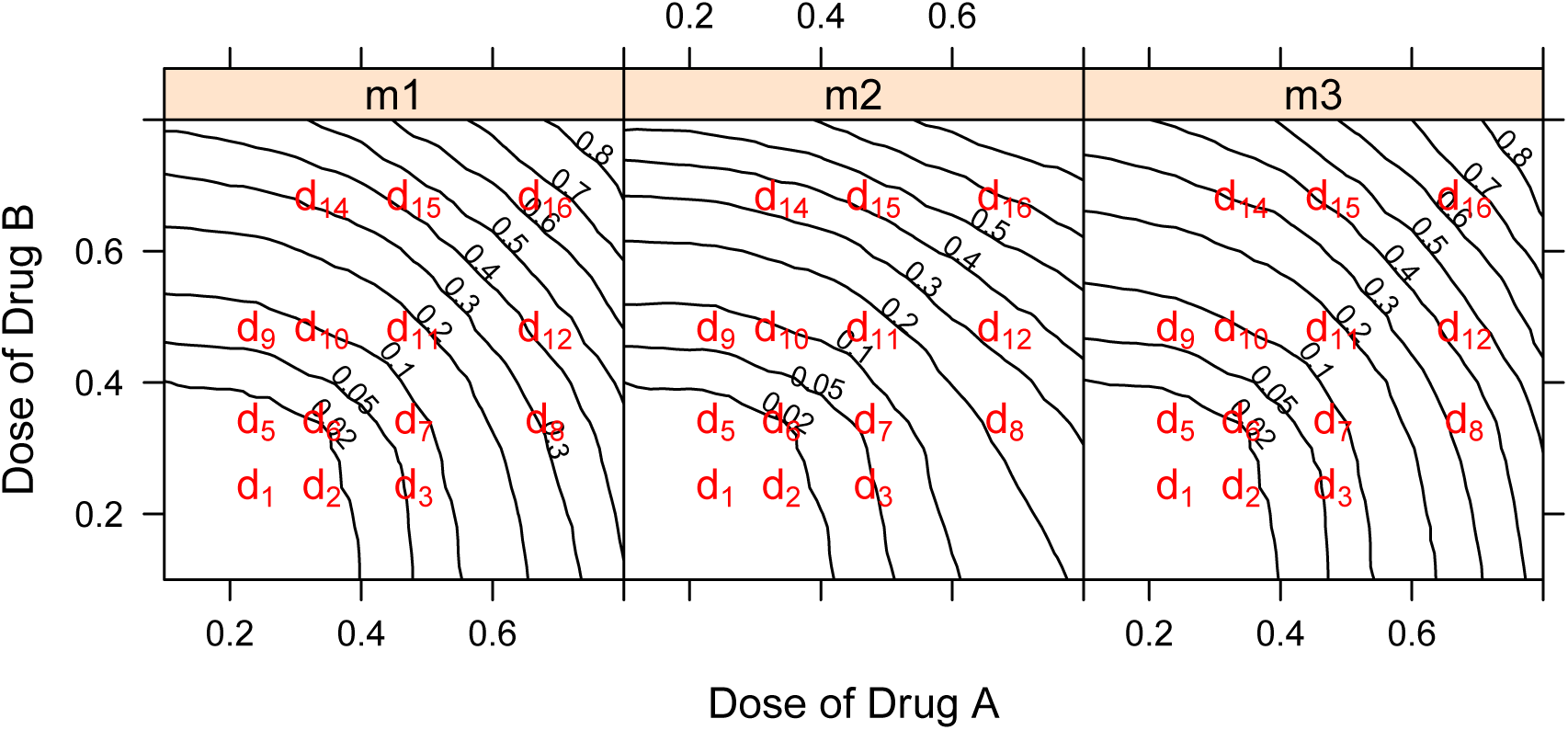
Obtaining alternative toxicity orderings by distorting the simulated DLT rates. The ‘true’ *m*_1_ model uses the rates calculated from our *N* = 1000 simulated patients (of whom the first 24 are shown in Figure 3). Alternative models *m*_2_ and *m*_3_ are obtained by gently distorting these DLT rates, as shown in the center and right panels.

**Table 2.** Three models used in POCRM, with DLT rates for the ‘true’ model *m*_1_.

|  | 1 | 2 | 3 | 4 | 5 | 6 | 7 | 8 | 9 | 10 | 11 | 12 | 13 | 14 |
| --- | --- | --- | --- | --- | --- | --- | --- | --- | --- | --- | --- | --- | --- | --- |
| $P_{DLT}$ | .003 | .007 | .014 | .021 | .061 | .073 | .083 | .100 | .178 | .311 | .333 | .432 | .451 | .665 |
| $\mathbf{m}_1$ | $d_1$ | $d_5$ | $d_2$ | $d_6$ | $d_3$ | $d_9$ | $d_7$ | $d_{10}$ | $d_{11}$ | $d_8$ | $d_{14}$ | $d_{12}$ | $d_{15}$ | $d_{16}$ |
| $\mathbf{m}_2$ | $d_1$ | $d_5$ | $d_2$ | $d_6$ | $d_3$ | $d_7$ | $d_9$ | $d_{10}$ | $d_{11}$ | $d_8$ | $d_{12}$ | $d_{14}$ | $d_{15}$ | $d_{16}$ |
| $\mathbf{m}_3$ | $d_1$ | $d_5$ | $d_2$ | $d_6$ | $d_3$ | $d_9$ | $d_7$ | $d_{10}$ | $d_{11}$ | $d_{14}$ | $d_8$ | $d_{15}$ | $d_{12}$ | $d_{16}$ |

<sup>4</sup>Notably, these subscripts are widely regarded as bad-luck numbers.

In this simulation, we find that 1-size-fits-all dosing at ‘the’ MTD selected by POCRM achieves a population-level remission rate with expectation 0.323. This represents a loss of 18.2% relative to the optimal 1-size-fits-all remission rate of 0.396. Considering that this reflects both the loss of ‘oracular’ knowledge about outcomes, and the coarse grid of available doses to search over, POCRM seems to have performed quite well *as a 1-size-fits-all dose-finding method* in this scenario.

## NAVIGATING 2-DIMENSIONAL TITRATION

Any realistic description of a dose-titration design requires a *graded* conception of DLT, enabling us to approach toxicity boundaries (such as the toxicity isocontours of Figure 3) without frequently crossing them. Because I have retained a binary (all-or-nothing) toxicity concept in this paper, I make no attempt here to specify a combination-dose titration algorithm explicitly.

Instead, I simply appeal to the characteristic toxicities of drugs A and B, together with the *dose-intensity* concept of (Simon and Korn 1990), as ‘navigational aids’ that make dose-titration in the (*a, b*)-plane conceivable. With synergy in mind, we posit a dose-intensity measure:

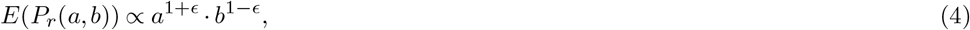

where ϵ = 0.01 serves simply to break the symmetry of our grid {*d*_1_, …*, d*_16_} about the 45° line. We may then ask what population remission rate is achieved by titrating each patient within the same grid used in the POCRM example, aiming to maximize (4) while preserving tolerability. As it turns out, this pragmatic dose titration over just 14 doses achieves a remission rate of 0.522. Relative to the remission rate of 0.586 attained by oracular dose individualization on a (near-)continuum of doses (*a, b*) ∈ [0, 2] *×* [0, 2], this represents an efficiency loss of just 11%.

## DISCUSSION

Figure 7 summarizes the performance results obtained here, placing the efficacy of two pragmatic methods (POCRM, titration) in context with corresponding ‘oracular’ methods. Clearly, in the scenario considere here, efficacy losses incurred in 1-size-fits-all dosing substantially exceed those due to pragmatism. Thus, considerations of pragmatism seem neither to justify the decision to use 1-size-fits-all dose-finding designs, nor to excuse their inefficiencies.

**Figure 7.**
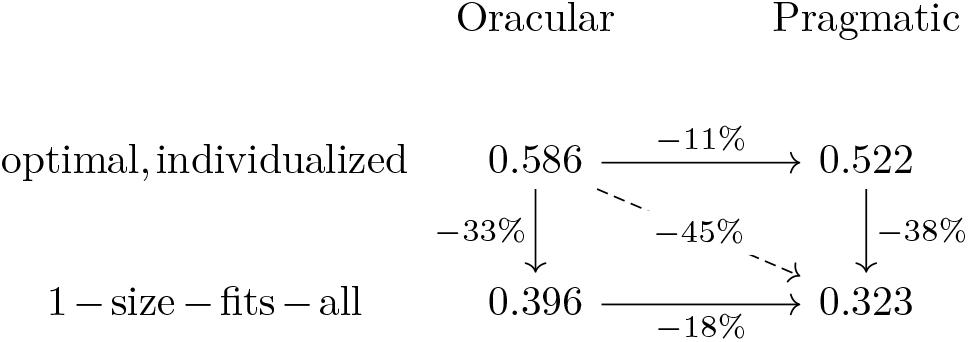
Efficacy losses attributable to several constraints, relative to ‘oracular’ optimal, individualized dosing on a continuum (top, left corner). In the specific scenario simulated in this paper, the 1-size-fits-all constraint (bottom row) appears substantially more costly than the ‘pragmatism’ of dosing on a coarse grid (right column). Remarkably, 1-size-fits-all dosing proves more costly here than the loss of ‘oracular’ knowledge about outcomes.

Of course, the results above derive from a single scenario. But to characterize the performance of POCRM (or any other 1-size-fits-all dose-finding method) across a full range of pharmacological scenarios would be as pointless as it is impossible. The essential point of my efforts here is to place POCRM and similar 1-size-fits-all dose-finding methods into a setting imbued with some relevant pharmacological realism. In this setting, the stark error of the past 30 years of ‘innovation’ in model-based 1-size-fits-all dose-finding methodology becomes readily apparent—at least to a person with clinical sensibilities, or sound pharmacological instincts.

At this point in my campaign against 1-size-fits-all dose finding, I feel less inclined to address my argument to the 1-size-fits-all dose-finding methodologists than to their clinical collaborators.^5^

To clinicians, the reality which Figure 3 palpably approximates must seem unrecognizable in the barren artifice of Table 2. Considering that the latter fully suffices to provide the inputs needed for a POCRM simulation, POCRM’s abnegation of clinical realism becomes evident. I continue to hope that self-respecting clinicians, who may have spent decades cultivating their clinical and pharmacological intuitions, will demand more respect from their trial-methodologist collaborators. On this point, one can hardly do better than to recall the long-neglected advice which (Sheiner 1991) offered under the heading ‘Reasserting Epistemologic Authority’:

> If I am right about what the illness is, then the cure is simple: clinicians must regain control over clinical trials to assure that the important questions are addressed. Here is my prescription for action:
>
>
> > “If you don’t understand it, don’t do it.”
>
> If a study design seems clinically inappropriate, wasteful, or unlike real medicine, the clinical scientist should ask why, and neither be satisfied nor acquiesce until it makes sense to him or her.
>
> — — Lewis B. Sheiner, MD (1940–2004)

## CONCLUSIONS

I have advanced a visual idiom for appreciating multiple, heterogeneous dose-limiting toxicities in the combination-therapy setting. In a simulation scenario, I have examined the population-level efficacy of two pragmatic dose-finding designs: a dose-titration method that maximizes dose-intensity subject to tolerability, and the well-known POCRM method for 1-size-fits-all combination-dose finding. Hypothetical ‘oracular’ methods serve to define theoretical upper limits of performance for individualized and 1-size-fits-all dosing, enabling separate consideration of the losses due to ‘pragmatism’ (i.e., the lack of ‘oracular’ knowledge of outcomes) and ‘1-size-fits-all-ness’. In the scenario considered here, 1-size-fits-all-ness makes a poor showing.

## DATA AVAILABILITY

### Open Science Framework

Code for reproducing all of this paper’s Figures and analyses may be found at doi:10.17605/osf.io/yn2q3.

## Footnotes

1 Consider the limiting case where A and B belong to the same drug class, or where a solvent used in both formulations causes the toxicity.

2 That is, I suppose the several toxicity-disutilities at any given dose combine via 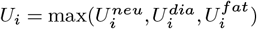, rather than additively or even multiplicatively as one might expect at least with simultaneious toixicitieis of comparable subjective severity. This assumption goes hand-in-hand, however, with the binary (all-or-nothing) toxicity conception which I employ throughout this paper, and which constitutes the true ‘original sin’ here. See the later section titled, ‘Navigating 2-Dimensional Titration’.

3 In (Norris 2017b) the 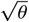 form of the dose-response was motivated as a means for obtaining an intermediate result in closed form. In the present paper, I pursue all results by strictly numerical means; the form of Equation 3 serves here simply to maximize continuity with the earlier argument.

4 Notably, these subscripts are widely regarded as bad-luck numbers.

5 Importantly, I take pains also to engage the patients who enroll in these trials, and their advocates. See e.g. https://www.growkudos.com/publications/10.1101%25252F240846/reader

